# PopMAG: A Nextflow pipeline for population genetics analysis based on Metagenome-Assembled Genomes

**DOI:** 10.64898/2026.01.21.700883

**Authors:** Daniel Sabogal-Rodriguez, Alejandro Caro-Quintero

## Abstract

Metagenome-assembled genomes (MAGs) are routinely recovered from metagenomic studies, yet the population genetic information embedded within these datasets remains largely underutilized. Analyzing within-species genetic variation can reveal adaptive evolution, selection pressures, and ecological dynamics that are hidden when MAGs are treated as homogeneous entities. Existing tools address individual analysis steps in isolation, requiring manual integration and creating barriers for researchers without extensive bioinformatics expertise. Here we present PopMAG, a Nextflow pipeline and interactive Shiny application that automates population genetics analysis of MAGs. PopMAG integrates quality control, community profiling, competitive read mapping, functional annotation, and microdiversity estimation into a single reproducible workflow. The pipeline calculates key population genetics metrics including nucleotide diversity (*π*), *pN/pS* ratios, fixation index (*F*_*ST*_), Levins’ index and SNVs counts with results consolidated into an interactive visualization platform for metadata-driven exploration. We demonstrate PopMAG’s utility through analysis of longitudinal cystic fibrosis lung metagenomes, where we detect signatures of antibiotic-driven selection in *Pseudomonas aeruginosa* efflux pump genes coinciding with treatment intervention.

**Availability and implementation:** PopMAG and corresponding documentation are publicly available at https://github.com/daasabogalro/PopMAG.

## 1 Introduction

Metagenomic studies routinely recover metagenome-assembled genomes (MAGs) to characterize the taxonomic and functional composition of microbial communities [Sun et al., 2025]. However, the population genetic information embedded within metagenomic datasets remains largely and with increasing interest of being characterized [Zhao et al., 2023]. A MAG represents a consensus sequence derived from multiple cells of closely related organisms, potentially distinct subpopulations or strains coexisting within a sample [Van Rossum et al., 2020]. This population structure possesses critical information about microbial adaptation, selection pressures, and ecological dynamics that is hidden when MAGs are treated as single, homogeneous entities.

Analysis of within-species genetic variation (microdiversity) in metagenomic data can reveal adaptive evolution in response to environmental change, host-microbe coevolution, antibiotic resistance emergence, and niche differentiation among co-occurring strains [Ghazi et al., 2022]. Population genetics metrics such as nucleotide diversity (*π*), fixation indices (*F*_*ST*_), and *pN/pS* ratios provide quantitative frameworks for detecting selection, inferring demographic history, and identifying genes under adaptive pressure. Despite the biological and clinical relevance of these insights, few metagenomic studies exploit the full population genetic potential of their data, often limiting analyses to MAG presence/absence or gene content comparisons.

Several bioinformatics tools enable population genetics analyses of metagenomic data, including inStrain [Olm et al., 2021], MetaPop [Gregory et al., 2022], POGENOM [Sjöqvist et al., 2021], Anvi’o [Eren et al., 201], M DAS [Nayfach et al., 201], and metaSNV [Costea et al., 2017]. However, these tools typically address individual steps in isolation, such as SNV calling, diversity estimation, or visualization, requiring users to manually integrate outputs, format data across platforms, and navigate complex installation dependencies. No existing solution provides a reproducible, end-to-end pipeline that combines MAG quality control, competitive read mapping, comprehensive population genetics analyses, and interactive visualization in a single automated workflow. This fragmentation creates barriers for researchers without extensive bioinformatics expertise and hinders reproducibility across studies.

Here we present PopMAG, an integrated scalable pipeline and Shiny application for population genetics analyses of MAGs that addresses these limitations. Our tool automates the complete workflow from quality-controlled MAGs and metagenomic reads to interactive visualizations, implementing competitive mapping strategies to ensure accurate strain-level resolution and providing interactive, metadata-driven exploration of evolutionary patterns. We demonstrate the pipeline’s utility through a case study of antibiotic-driven selection in *Pseudomonas aeruginosa* populations from a cystic fibrosis patient, revealing adaptive evolution in a multidrug efflux pump gene coinciding with treatment intervention.

## 2 Implementation

We developed a Nextflow [Di Tommaso et al., 2017] pipeline for population genetics analysis of metagenome assembled-genomes (MAGs) that integrates quality control, competitive mapping, functional annotation, and microdiversity estimation into a single automated workflow. The pipeline accepts three primary inputs: (1) a samplesheet of assembled MAGs, (2) a samplesheet of metagenomic reads, and (3) a metadata file for downstream visualization.

The workflow consists of five major stages (Fig. 1). First, MAG quality control assesses genome completeness and contamination using CheckM2 [Chklovski et al., 2023], filters bins based on user-defined quality thresholds, and performs dereplication with dRep [Olm et al., 2017] to generate a non-redundant genome database. Second, community profiling employs SingleM [Woodcroft et al., 202] to characterize the taxonomic composition and provide context for population-level analyses. Third, competitive mapping creates a dereplicated genome database, generates contig-to-bin files, performs competitive mapping with Bowtie2 [Langmead and Salzberg, 2012], and calculates per MAG coverage using CoverM [Aroney et al., 2025]. Fourth, annotation and microdiversity analysis uses Prodigal [Hyatt et al., 2010] for gene prediction, MetaCerberus [Figueroa III et al., 2024] for functional annotation, inStrain [Olm et al., 2021] for within-species diversity profiling (including nucleotide diversity *π* and *pN/pS* ratios), and POGENOM [Sjöqvist et al., 2021] for between-population differentiation (*FST*). Finally, visualization consolidates in-Strain profiles, POGENOM results, annotation summaries, and coverage statistics into formatted outputs that feed directly into an interactive Shiny application.

**Figure 1.**
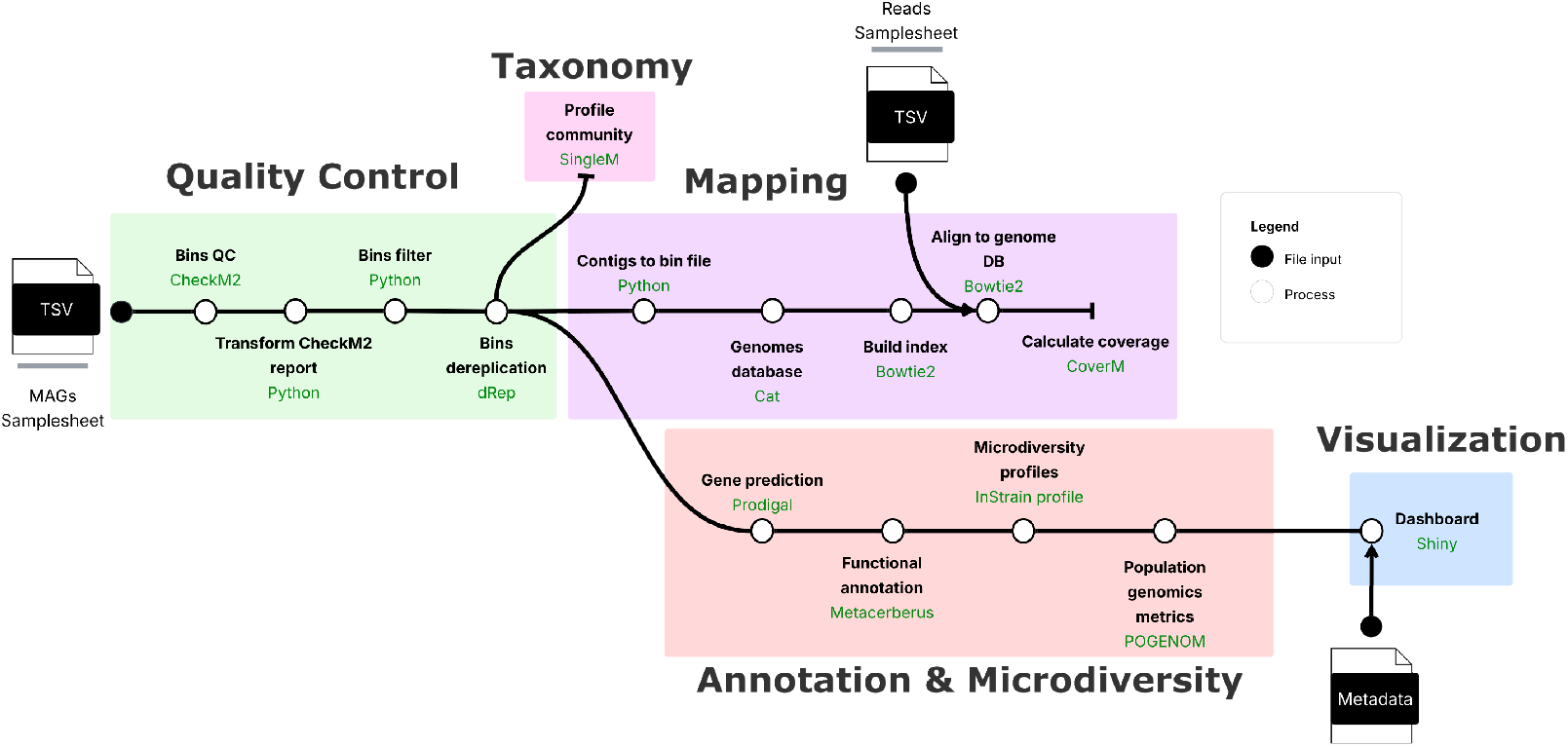
Workflow overview. Schematic representation of PopMAG. The workflow consists of five major stages: (1) MAG quality control, (2) community profiling, (3) competitive read mapping, (4) functional annotation and microdiversity analyses, and (5) data integration and visualization through an interactive Shiny application. Each one of the five steps is represented by a different colour. The required input files, including the metadata file, are represented as filled circles, while the different processes are represented as empty circles.

The pipeline employs containerization (Docker/Singularity) [Merkel, 2014, Kurtzer et al., 2017] for reproducibility and implements parallel processing of samples to ensure scalability across datasets of varying sizes. Unlike existing tools that address individual steps in isolation, our end-to-end implementation eliminates the need for manual data formatting and transfer between analysis stages. The integrated Shiny application enables interactive exploration of results through metadata-driven visualizations, allowing researchers to dynamically filter, compare, and export population genetics metrics across experimental conditions or sample groups. All intermediate and final outputs are organized in a standardized directory structure for easy interpretation and downstream analysis.

### 2.1 Technical architecture

The pipeline is implemented in Nextflow (version ≥ 24.10.0) using DSL2 syntax for modular process design. All bioinformatics tools are packaged in individual containers available via Quay.io, with support for both Docker and Singularity execution. Alternative dependency management through Conda/Mamba environments is available for users preferring non-containerized execution.

The pipeline implements a six-level resource labelling system ranging from lightweight single-CPU processes (6 GB RAM) to high-memory operations (200 GB RAM), with automatic error recovery through dynamic resource scaling, failed processes retry with doubled CPU and memory allocations. Pre-configured execution profiles support local workstations, SLURM-managed HPC clusters, and AWS cloud infrastructure. Users customize pipeline behaviour through a documented configuration file or command-line parameters.

Nextflow automatically parallelizes processes across samples and within-sample per-MAG analyses (annotation, diversity profiling) based on available resources and workflow dependencies, enabling efficient scaling from small studies to large metagenomic surveys.

### 2.2 Core functionalities

#### MAG Quality Control and Dereplication

The pipeline filters MAGs using stringent quality thresholds (≥ 90% completeness, ≤5% contamination) as assessed by CheckM2, ensuring high-confidence genomes for population-level inference. These thresholds con be configured by the user to accommodate different study requirements. dRep performs genome dereplication using default parameters to generate a non-redundant genome set, eliminating near-identical genomes that would confound competitive mapping.

#### Competitive Mapping and Coverage Estimation

To minimize read mis-mapping in complex metagenomic samples, the pipeline implements competitive read alignment where reads map simultaneously against all dereplicated genomes. This approach ensures that each genome in the database is sufficiently distinct to avoid mapping confusion and significantly reduces false-positive microdiversity signals. Bowtie2 performs the competitive alignment, and CoverM calculates genome coverage using trimmed mean (10-90th percentile) with a minimum covered fraction threshold of 0.5, providing robust coverage estimates that are resilient to coverage outliers.

#### Microdiversity and Population Genetics Analyses

The pipeline integrates complementary tools for comprehensive evolutionary inference. inStrain profiles within-population diversity, calculating nucleotide diversity (*π*), *pN/pS* ratios, linkage patterns, and SNV characteristics using default quality filters. POGENOM performs between-population analyses, computing *π* within each sample and pairwise *F*_*ST*_ values across all sample pairs when multiple populations are available. When genomic annotations are provided, POGENOM calculates gene-wise *π* and *F*_*ST*_ at both nucleotide and amino acid levels, enabling detection of selection at the gene level. The pipeline also computes Levins’ niche breadth index using the microniche R package, quantifying the ecological specialization of each population across samples.

#### Functional Annotation

Prodigal predicts protein-coding genes, which are functionally annotated by MetaCerberus against comprehensive databases including KEGG (KO), COGs, CAZy, FOAM, VOGs, and PHROGs. Annotation results are integrated with microdiversity metrics, enabling users to link evolutionary patterns (*pN/pS* ratios, gene-wise *F*_*ST*_) to functional categories and metabolic pathways.

#### Data Integration

All outputs, inStrain profiles, POGENOM statistics, functional annotations, and coverage data are automatically merged into standardized tab-separated value (TSV) tables formatted for direct import into the Shiny app visualization, eliminating manual data wrangling steps.

### 2.3 Shiny application

The pipeline automatically launches an interactive Shiny application for exploratory data analysis and visualization of population genetics results. The application is configured directly from the pipeline, with customizable port settings and session duration to accommodate different deployment scenarios (local workstations, shared servers, or cloud instances).

The Shiny App features a sidebar for MAG selection and dynamic plot customization, allowing users to adjust visual parameters (colours, point sizes, axis scales) in real-time. A top navigation bar organizes visualizations into two main modules: metadata-driven analyses and MAG-specific population genetics metrics. The metadata section enables cross-sample comparisons through a correlation heatmap visualizing relationships between environmental or experimental variables, and principal component analysis (PCA) for dimensionality reduction and sample clustering. Users can interactively filter samples based on metadata criteria to explore condition-specific patterns. For each selected genome, the application provides five complementary visualizations: (1) Levins’ niche breadth index indicating ecological specialization across samples, (2) nucleotide diversity (*π*) barplots comparing within-population variation, (3) scatter plots relating nucleotide diversity (*π*) to individual metadata variables with metadata-based filtering, (4) SNVs and SNSs count distributions across samples, and (5) histograms of gene-wise *pN/pS* ratios with an interactive, filterable table displaying gene annotations and corresponding selection metrics (Supplementary data). This integrated view enables users to identify genes under selection and their functional roles.

Each visualization is accompanied by its source data table, providing transparency and enabling users to inspect underlying values. Users can download plots and use filtered data tables for custom downstream analyses or reporting. The integration between pipeline outputs and visualization ensures that results are immediately accessible through an intuitive interface, lowering barriers for researchers without extensive bioinformatics expertise.

## 3 Case study

To demonstrate the pipeline’s utility for detecting adaptive evolution in clinical metagenomes, we analysed an open access longitudinal data set of sputum samples from a cystic fibrosis (CF) patient undergoing antibiotic therapy. The dataset consisted of 10 metagenomic samples collected over 19 months, with sequencing depths ranging from 11.9 to 43.6 million human-filtered reads per sample. The patient received continuous treatment with azithromycin, colistin, and insulin throughout the study period, with the CFTR modulator lumacaftor/ivacaftor added to the regimen at month 13 [Dmitrijeva et al., 2021].

The complete workflow processed all 10 samples in 8.5 hours on a local workstation with 12 CPUs and 128 Gb of RAM, with inStrain profiling accounting for approximately 5.6 hours of runtime. BAM files ranged from 300 MB to 3.3 GB in size. Quality control identified six high-quality MAGs (≥ 90% completeness, ≤5% contamination) representing key CF associated taxa: *Pseudomonas aeruginosa, Achromobacter sp*., *Fusobacterium sp*., *Parvimonas sp*., *Prevotella sp*., and *Streptococcus sp*. All six MAGs passed quality filters and were included in downstream analyses.

Using the Shiny application’s filterable and interactive gene-level *pN/pS* table, we identified a multidrug efflux pump subunit (MexA, membrane-fusion protein) in the *P. aeraginosa* MAG (mean coverage 155X) exhibiting a selection signature specifically in the post lumacaftor/ivacaftor treatment sample (pN/pS = 2.1). In the nine pre-treatment samples spanning months 0-13, this gene showed no evidence of positive selection. The elevated pN/pS ratio indicates positive selection for nonsynonymous mutations in MexA coinciding with lumacaftor/ivacaftor administration, suggesting adaptive resistance evolution. Notably, this gene was not previously reported among mutations associated with CFTR modulator treatment [Ledger et al., 2024] which identified mutations in genes such as *mutS, mutL, algG*, and *mucA*.

Additional genes in *P. aeraginosa* displayed signatures of positive selection at various timepoints, including an AlgP family protein (pN/pS = 4.3), Cbb3-type cytochrome oxidase subunit 1 (2.5), fibronectin-binding autotransporter adhesin (2.2), and nitrate/nitrite transporter NarK (2.0), revealing temporally dynamic evolutionary responses throughout the treatment course.

Integration of clinical metadata revealed an inverse relationship between *P. aeraginosa* nucleotide diversity (*π*) and C-reactive protein (CRP) concentration, with microdiversity decreasing as inflammation markers increased. This pattern, readily visualized through the Shiny app’s metadata-linked diversity plots (Fig. 2), suggests that host immune responses or treatment-associated pressures constrain within-population genetic variation during inflammatory episodes.

**Figure 2.**
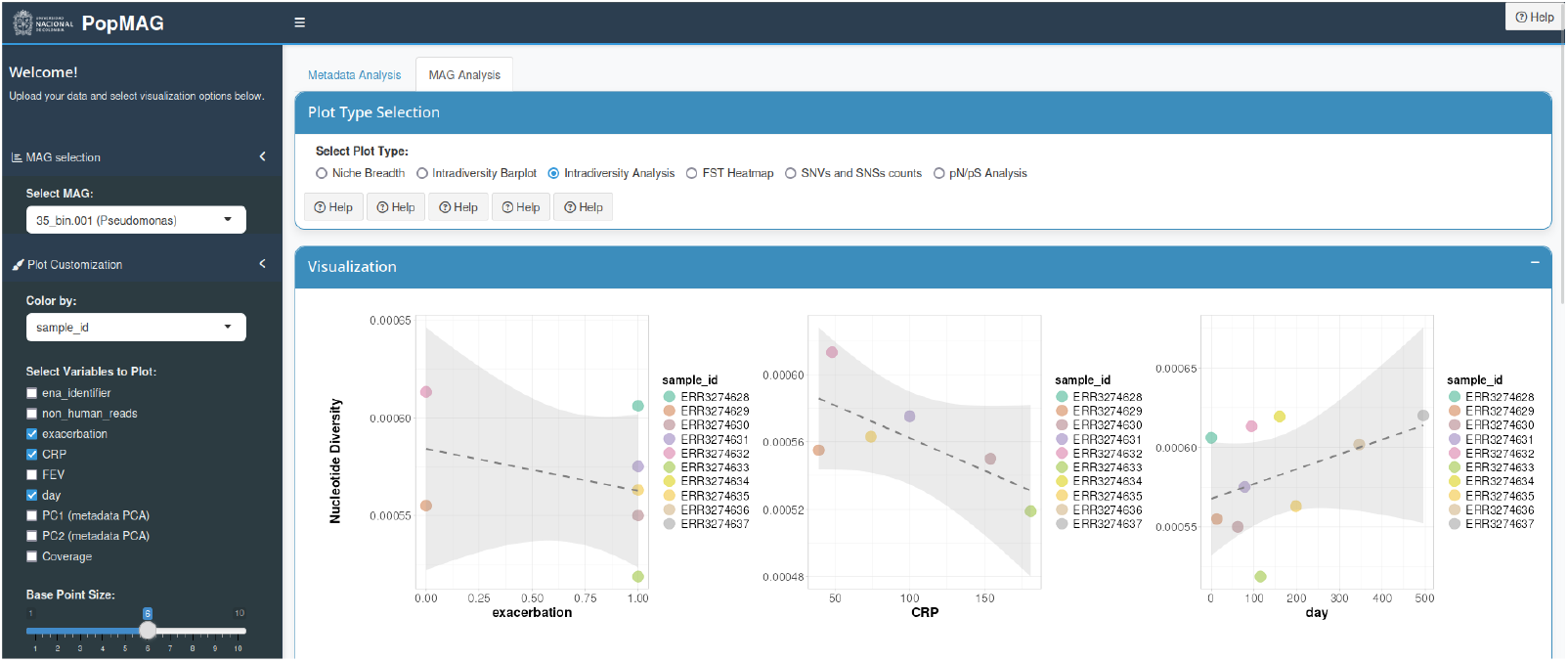
Interactive Shiny application interface for population genetics exploration. Structure of the shiny app generated after the PopMAG workflow execution. The MAG to be analysed can be selected using the sidebar on the left and the different plot types can be selected in the navigation bar at the top. The sidebar also features a “Plot Customization” section to configure the visualizations. In this example we have selected the *P. aeraginosa* MAG and we’re visualizing the “Intradiversity Analysis” plots, where we can see the relationship between different metadata values (exacerbation, CRP and day, as selected in the sidebar) and the nucleotide diversity (*π*).

This case study demonstrates the pipeline’s capability to identify relevant adaptive evolution in pathogen populations, link selection signatures to treatment interventions through metadata integration, and show biologically interpretable patterns through intuitive visualizations, all within a single automated workflow.

## 4 Conclusions

We present an end-to-end Nextflow pipeline and interactive Shiny application for population genetics analyses of metagenome-assembled genomes, addressing the growing need for integrated tools to study microbial evolution in complex communities. By combining MAG quality control, competitive read mapping, microdiversity profiling, and functional annotation in a single automated workflow, our pipeline eliminates manual data formatting and streamlines the path from raw metagenomic data to interpretable evolutionary insights.

The pipeline’s modular architecture and containerized implementation ensure reproducibility and portability across computing environments. The integrated Shiny application democratizes access to population genetics metrics from MAGs through metadata-driven visualizations, allowing researchers without extensive bioinformatics expertise to explore adaptive evolution, ecological specialization, and selection signatures in their metagenomic data.

Our case study of *Pseudomonas aeruginosa* during cystic fibrosis treatment demonstrates the pipeline’s power to detect clinically relevant adaptive responses and link selection signatures to treatment interventions. The identification of antibiotic-driven selection in a multidrug efflux pump gene highlights how this approach can reveal novel resistance mechanisms that complement traditional genomic surveillance efforts.

The pipeline and Shiny application are freely available at https://github.com/daasabogalro/popmag, along with documentation, to facilitate adoption by the microbiome research community. This tool can contribute to accelerate research into microbial adaptation and the ecological and evolutionary dynamics shaping microbiome function.

## Conpeting interests

No competing interest is declared.

## Acknowledgnents

Funding was provided by the Universidad Nacional de Colombia and MINCIENCIAS through a grant awarded under the Max Planck Tandem Group initiative.

## Author contributions statenent

D. S-R developed the software, conducted formal analysis and validation, performed data curation and visualization, designed the methodology, drafted the original manuscript, and approved the final version. A.C-Q contributed to conceptualization and methodology, conducted formal analysis, provided resources and funding acquisition, supervised the project, contributed to manuscript review and editing, and approved the final version.

