## supplementary file 1 for "PopMAG: A Nextflow pipeline for population genetics analysis based on Metagenome-Assembled Genomes"

[1] Departamento de Ingeniería de Sistemas e Industrial, Universidad Nacional de Colombia, 111321, Bogotá, Colombia.

[2] Departamento de Biología, Facultad de Ciencias, Universidad Nacional de Colombia, 111321, Bogotá, Colombia.

[3] Max Planck Tandem Group in Holobionts, Universidad Nacional de Colombia, 111321, Bogotá, Colombia.

Supplementary figure S1: Bar chart displaying coverage distribution of MAG 35_bin.001 across 10 longitudinal cystic fibrosis lung metagenome samples, with proportional abundances shown above bars. The normalized Levins' index (Bn = 0.617) indicates a moderately generalist distribution, with the MAG present in all samples but at varying abundances. Samples are ordered by decreasing coverage.

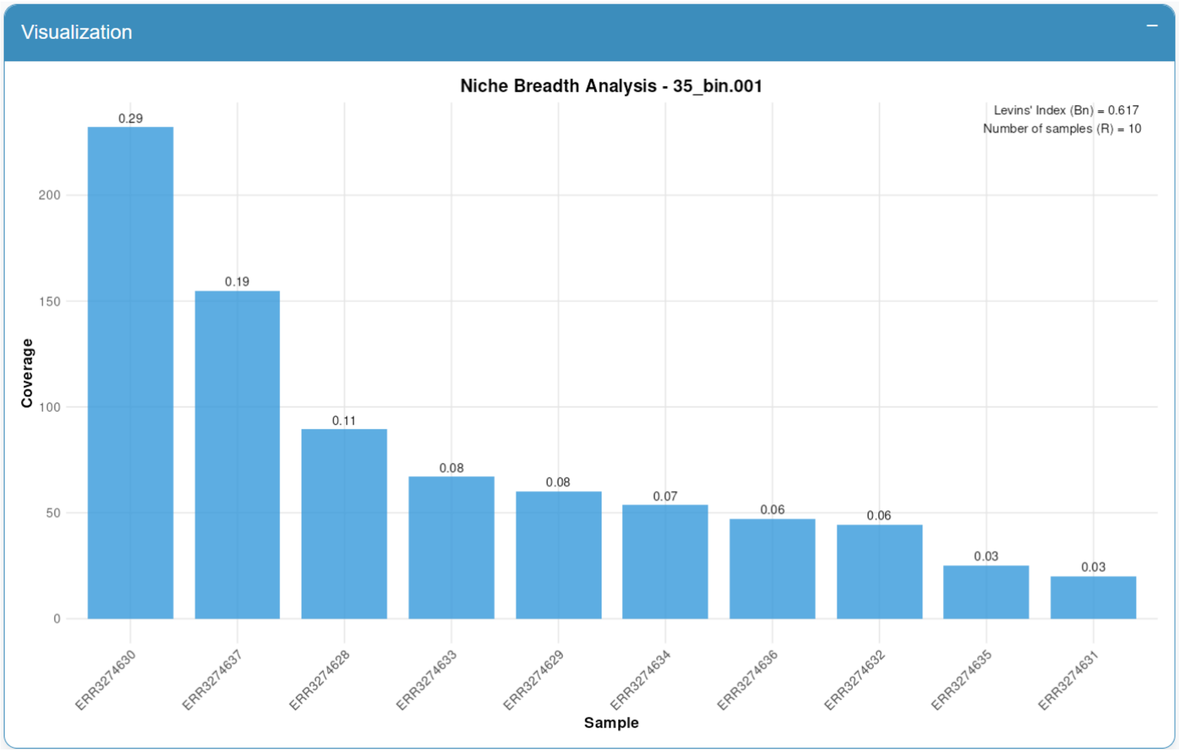

Supplementary figure S2: Bar chart displaying nucleotide diversity (π) for MAG 35_bin.001 across 10 longitudinal cystic fibrosis samples. The red line indicates 50× rarefied diversity to account for coverage variation. A marked decrease in diversity at sample ERR3274631 suggests a potential population bottleneck or selective event. Diversity values range from ~$7 \times 10^{-5}$ to ~$6 \times 10^{-4}$.

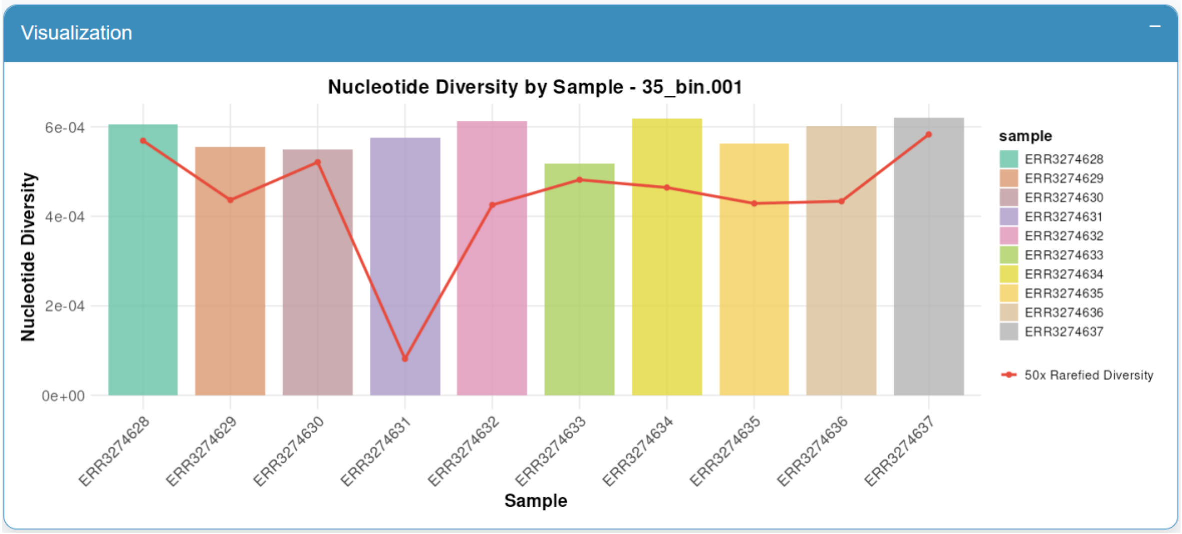

Supplementary figure S3: Scatter plots correlating nucleotide diversity (π) with clinical metadata: exacerbation status (left), C-reactive protein levels (center), and sampling day (right). Points are colored by sample ID and sized by coverage. Dashed lines show linear regression fits with 95% confidence intervals. This metadata-driven visualization enables exploration of associations between population genetic metrics and clinical covariates.

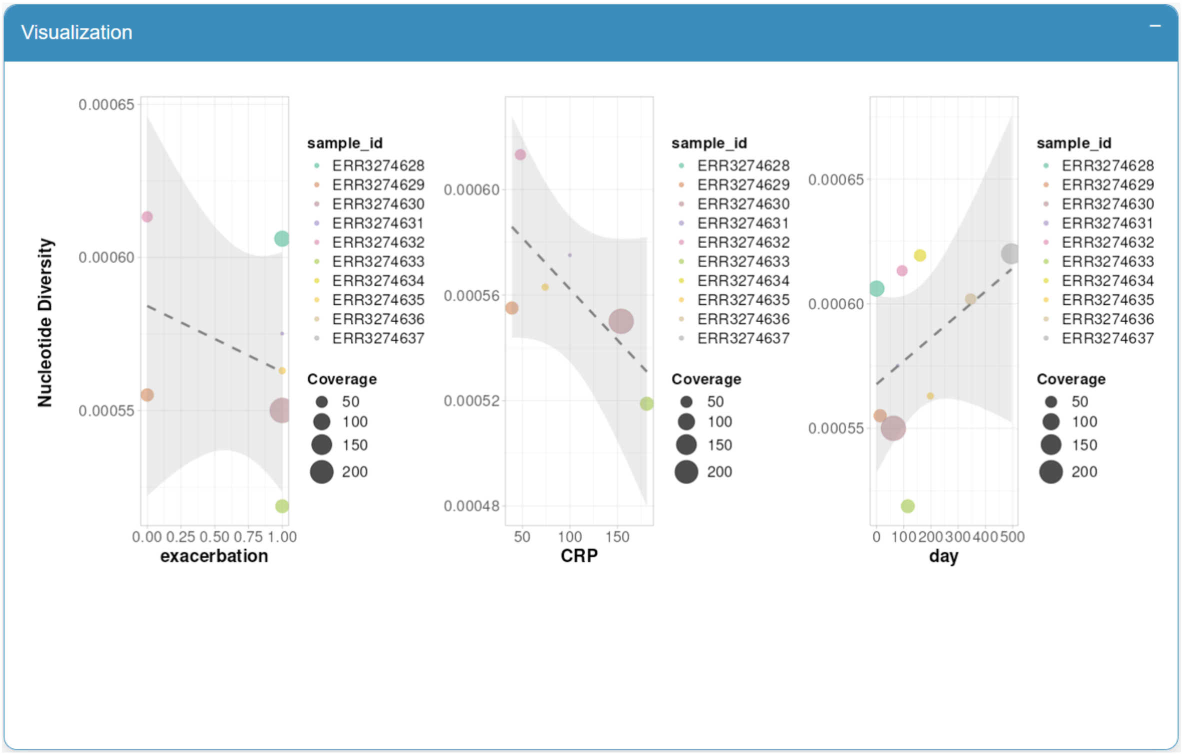

Supplementary figure S4: Heatmap displaying fixation indices (FST) between all sample pairs for MAG 35_bin.001 across longitudinal cystic fibrosis samples. Color scale ranges from yellow (FST ≈ 0, low differentiation) to red (FST ≈ 0.5, high differentiation). Elevated FST values involving ERR3274630 (up to 0.307) indicate population structure changes at this time point, potentially reflecting strain replacement or selective sweeps.

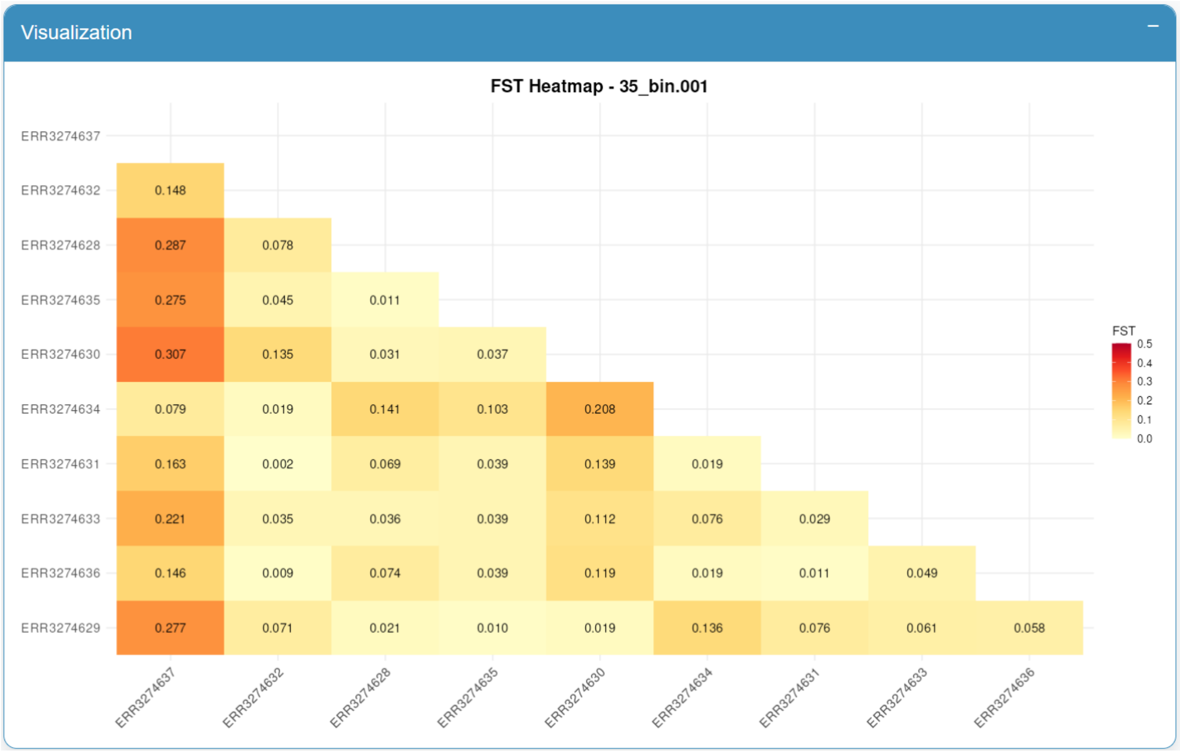

Supplementary figure S5: Histogram showing the distribution of gene-level pN/pS ratios for MAG 35_bin.001 across all samples. The dashed red line indicates neutral selection (pN/pS = 1). The strong left-skew (majority of genes with pN/pS < 0.1) indicates widespread purifying selection, while rare genes with pN/pS > 1 represent candidates under positive selection.

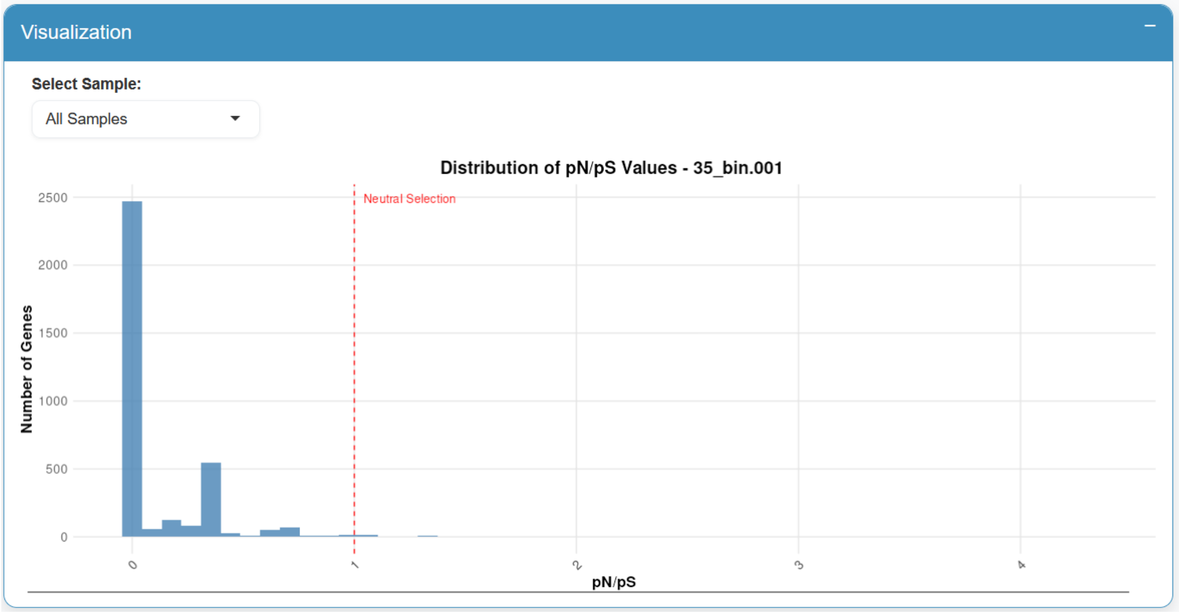

Supplementary figure S6: Interactive table displaying per-gene population genetics metrics including pN/pS, nucleotide diversity, coverage, and functional annotations. The search and filter functionality enables targeted exploration of genes of interest. Here, the MexA (AcrA) efflux pump gene shows elevated pN/pS (2.1) in sample ERR3274637 compared to earlier time points (pN/pS = 0.0), indicating positive selection potentially driven by antibiotic pressure.

Data Table

Search:

| Sample | Product | pN/pS | Nucleotide Diversity | Coverage | Breadth | Best Hit | EC Number |
|---|---|---|---|---|---|---|---|
| All | iit MexA | All | All | All | All | All | All |
| ERR3274637 | multidrug efflux RND transporter periplasmic adaptor subunit MexA | 2.1 | 0.0 | 162.6 | 1.0 | NF033834.1 | |
| ERR3274628 | multidrug efflux RND transporter periplasmic adaptor subunit MexA | | 0.0 | 109.3 | 1.0 | NF033834.1 | |
| ERR3274629 | multidrug efflux RND transporter periplasmic adaptor subunit MexA | | 0.0 | 74.4 | 1.0 | NF033834.1 | |
| ERR3274630 | multidrug efflux RND transporter periplasmic adaptor subunit MexA | | 0.0 | 266.3 | 1.0 | NF033834.1 | |
